# Retroelement expansions underlie genome evolution in stingless bees

**DOI:** 10.1101/2025.04.12.648530

**Authors:** Natalia de Souza Araujo, Patricia Azevedo, Rafael Rodrigues Ferrari, Lucas Borges dos Santos, Florence Rodriguez, Roberta Cornélio Ferreira Nocelli, Matthew Hudson, Thiago Mafra Batista, Klaus Hartfelder, Serge Aron

## Abstract

Stingless bees are essential pollinators and emerging models for studying behavioral and genomic evolution. In the genus *Melipona*, a major difference in heterochromatin organization defines two groups: Group I species (e.g., *M. quadrifasciata*) with <50% of pericentromeric heterochromatin and Group II species (e.g., *M. scutellaris*) containing >50% heterochromatin across their chromosomes. These differences are thought to correlate with genome size and transposable element (TE) content, offering a unique opportunity to explore how heterochromatin variation, TE dynamics, and chromosomal evolution interact in a phylogenetic context. We present pseudo-chromosome-level genome assemblies for *M. quadrifasciata* and *M. scutellaris* obtained by long-read sequencing and Hi-C scaffolding. Comparative analyses reveal conserved synteny but marked divergence in structural variants and TEs. *M. scutellaris* shows an expansion of retrotransposons, particularly Gypsy/DIRS1 elements, concentrated in TE hotspots linked to chromosomal rearrangements and structural variants. This coincides with distinct methylation entropy and an expansion of histone deacetylase orthologs, potentially affecting heterochromatin organization. The increased ratio of retrotransposons in *M. scutellaris* is counterbalanced by more DNA transposons in *M. quadrifasciata*, resulting in genomes of similar overall sizes but of distinct heterochromatin distribution. Advancing our understanding of genome evolution in eusocial insects, we provide high-resolution genomic resources for two *Melipona* species that differ in heterochromatin content. Our results highlight the complex role of TEs in shaping genomes and underscore their influence on chromosomal and epigenetic innovation, providing strong evidence that TE dynamics underly the striking heterochromatic differences observed in *Melipona*.

## Introduction

Bees are critical pollinators of both natural ecosystems and crops, playing a key role in global food security and environmental stability (Ollerton et al. 2011; Potts et al. 2016). Native bee species, in particular, foster sustainable ecosystems by preserving native flora diversity while enhancing agricultural yields (Jaffé et al. 2016; Bueno et al. 2023). With over 20,000 species described worldwide, bees represent a highly diverse group of insects that exhibit a wide range of ecological strategies and behaviors, many of which remain poorly understood (Michener 2007; Almeida et al. 2023). Native to tropical and austral subtropical regions, stingless bees (Hymenoptera: Apoidea: Anthophila: Meliponini) encompass nearly 600 species across 60 genera (Engel et al. 2023; Michener 2007) that humans have managed since at least 300 BC (Hrncir et al. 2016). Like honeybees, stingless bees are eusocial, forming large colonies with a caste-based division of labor, in which queens specialize in reproduction and a multitude of female workers are dedicated to various colony functions, such as foraging, colony maintenance, and defense (Hrncir et al. 2016; Michener 2007; Engel et al. 2023). Stingless bees are remarkably versatile pollinators, visiting an array of 1,435 different plant genera from 215 families (82 families in the Afrotropical region, 137 in the Indo-Malayan-Australasian region, and 205 in the Neotropics) (Bueno et al. 2023). The rich species diversity in this group is reflected in their morphological, ecological, and behavioral traits (Michener 2007; Grüter 2020; Bueno et al. 2023; Hrncir et al. 2016). Despite their environmental relevance, significant gaps remain in our understanding of stingless bee biology and genome evolution, as only 14 stingless bee genomes (∼2.8% of the species) have been fully sequenced to date, none achieving chromosome-level resolution (2024).

Eusocial bee genomes are notable for their exceptionally high recombination rates, reaching up to 19 cM/Mbp in the honeybee (*Apis mellifera*) (Beye et al. 2006; Fouks et al. 2024). This elevated recombination rate is thought to have facilitated the evolution of eusociality (Ross et al. 2015; Wilfert et al. 2007). High recombination rates are typically associated with G/C-biased gene conversion (Kent and Zayed 2013; Chen et al. 2007). Paradoxically, eusocial bee genomes are overall highly A/T biased, with G/C-rich regions only scattered across the genome (Kent and Zayed 2013; Wallberg et al. 2015). Fine-scale recombination analyses of honeybee genomes suggest that this pattern may result from two distinct DNA double-strand break repair mechanisms in bees: one that leads to A/T enrichment and another consistent with classical G/C-biased gene conversions (Fouks et al. 2024). Thus, high recombination rates associated with G/C-biased gene conversion would have been selected to counteract A/T-rich genomes, where double-strand breaks with A/T-biased resolution frequently occur (Fouks et al. 2024).

Although the mechanisms driving recombination events are less well understood in eusocial bee lineages beyond *A. mellifera*, genomic recombination followed by chromosomal rearrangements likely contributes to chromosome number variation across corbiculate bees (i.e., the Euglossini [n=20-21], the Apini [n=17], the Bombini [n=18-20], and the Meliponini [n=7-18]) (Travenzoli et al. 2019; Cunha et al. 2021). Bees of the genus *Melipona* almost invariably possess nine chromosomes (Travenzoli et al. 2019). Still, some species exhibit substantial chromosomal variation characterized by distinct heterochromatic patterns (Pompolo et al. 2002; Pereira et al. 2021; Tavares et al. 2023). According to cytogenetic observations, *Melipona* species can be divided into two main groups based on their constitutive heterochromatin levels: Group I includes species in which less than half of the genome is composed of heterochromatin (primarily found in the pericentromeric region), and Group II gathers species with over 50% of heterochromatin in their genomes dispersed throughout the chromosomes (da Cunha et al. 2018; Pereira et al. 2021). Additionally, genome size, estimated through flow cytometry, differs between the two groups, with Group II species showing much larger genome size estimations (Tavares et al. 2010). Low-coverage short-read sequencing of Group I (*Melipona quadrifasciata*, subgenus *Melipona sensu stricto*) and Group II (*Melipona scutellaris*, subgenus *Michmelia*) species further suggests significant differences in repetitive element content (Pereira et al. 2021). While for *M. scutellaris,* over 50% of the data generated consisted of repeats, only 3.78% of *M. quadrifasciata* data was repetitive (Pereira et al. 2021). These findings suggest that a large quantitative difference in repetitive transposable elements (TEs) might exist between Group I and Group II species, possibly explaining the observed variation in heterochromatin patterns (Pereira et al. 2021; Tavares et al. 2023; Piccoli et al. 2018). Given the unique evolutionary pressures shaping eusocial bee genomes – such as high recombination levels and structural chromosomal differences – they serve as an excellent model for investigating the mechanisms driving rapid heterochromatin variation and chromosomal evolution within a phylogenetic frame.

We present a detailed genomic investigation of the differences between *Melipona* species of Group I and Group II to elucidate the mechanisms underlying their heterochromatin variation. We used *M. quadrifasciata* and *M. scutellaris* as representative species from Group I and Group II. These species diverged ∼14–17 million years ago (Ramírez et al. 2010), sharing the same chromosome number (n = 9) but differing in expected genome sizes and repetitive content (Pereira et al. 2021; Tavares et al. 2010). We assembled high-quality genomes for both species using long-read sequencing and DNA conformation data to produce scaffolds consistent with pseudo-chromosome molecules. Then, we conducted genomic comparative analyses, including genome synteny, gene and repetitive element content assessment, and structural variation analysis based on pangenome graphic tools. Our results reveal that genome synteny and gene content are largely conserved between species, but contrary to our expectations, their repetitive element content and genome sizes were also very similar. Our data shows that the major differences between these bees can be explained as arising from the nature of the TEs in their genomes. In *M. scutellaris* (Group II), retroelements have expanded, occupying positions containing DNA transposons in *M. quadrifasciata*, this likely caused numerous structural variants (SVs) between genomes and the observed heterochromatin differences without significantly affecting genome size.

## Results

### An improved genome assembly for *Melipona quadrifasciata*, the first complete pseudo-chromosome-level assembly of a stingless bee

Before scaffolding with conformation data, our *M. quadrifasciata* genome assembly comprised 85 contigs totaling around 256.14 Mb. The largest contig was over 33 Mb long, and the N50 and L50 values were 16 Mb and 7, respectively. No potential contaminant sequences were identified in this assembly (Figure S1). Of the 256.14 Mb of genomic data, 250,752,974 bp matched sequences from species of the family Apidae, with 4,398,558 bp matching species of the family Andrenidae, and smaller hits of 967,436 bp and 3,274 bp matching to Halictidae and Megachilidae, respectively. Nineteen contigs showed no significant matches (totaling 20,932 bp of unidentified data) and had notably low GC content (Figure S1). We identified 935,870 possible restriction sites (GATC) within the genome of *M. quadrifasciata* and generated 133,082,626 paired reads from the Hi-C library. Following alignment-based quality trimming, 74,945,927 paired reads (i.e., ∼41x genome coverage) were retained for the Hi-C matrix construction. Hi-C scaffolding, along with manual curation and correction of one translocation, reduced the assembly to 66 scaffolds, including one scaffold for the mitochondrial genome. Long reads were aligned back to the genome for a final quality assessment, yielding an estimated mean coverage of 88x (SD=573) with a mean mapping quality of 54.82. One scaffold (scaffold_66, length 89 bp) had no read alignments and was subsequently removed along with the mitochondrial contig.

The final nuclear genome of *M. quadrifasciata* consists of 64 scaffolds totaling 256,145,285 bp, with a mean GC content of 38.77% (Figure S2), an L90 on the eighth scaffold, and an N50 of 30,037,562 bp – almost as large as the maximum scaffold size of 34,390,260 bp (Figure 1, Table I). The Hi-C contact map confirms that the largest nine scaffolds represent nine pseudo-chromosomes, ranging from 34,390,260 bp to 23,026,935 bp (Figure 1), and contain 99,8% of the genome assembly (Figure S2). The BUSCO completeness score of the full genome assembly is 97.7% (Figure 1). Additionally, we identified 15 possible NUMT regions across eight scaffolds, ranging from 144 to 4,929 bp (Figure S3). A total of 26 ribosomal RNA (rRNA) genes (23 complete and three partial sequences) were identified (Table I). Genome annotation resulted in the prediction of 15,741 protein-coding genes, with a BUSCO completeness of 98.3% (C=98.3% [S=68.4%, D=29.9%], F=0.3%, M=1.4%), and 13,946 transcripts, with a mono-to-multi-exonic transcript ratio of 0.12 (monoexonic transcripts = 1,531 and multiexonic transcripts = 12,415). Of all the predicted genes, 14,445 (91.77%) matched database entries for functional annotation, including 12,484 hits against Pfam. Methylation was estimated for cytosines as regular methylations (Cm) and hydroxymethylation (Ch) from an estimated threshold above or equal to 0.78125 and for adenosines (A) with a threshold of 0.8027344. Mean Cm and Ch methylation rates in the genome were 0.96% and 0.27%, respectively, while mean A methylation was 1.4%. The distribution of the calculated methylation entropy of CpG sites based on 50bp windows across the genome is shown in Figure 2. A spread entropy distribution is observed in *M. quadrifasciata* as it would be expected from a whole-body sample.

**Figure 1.**
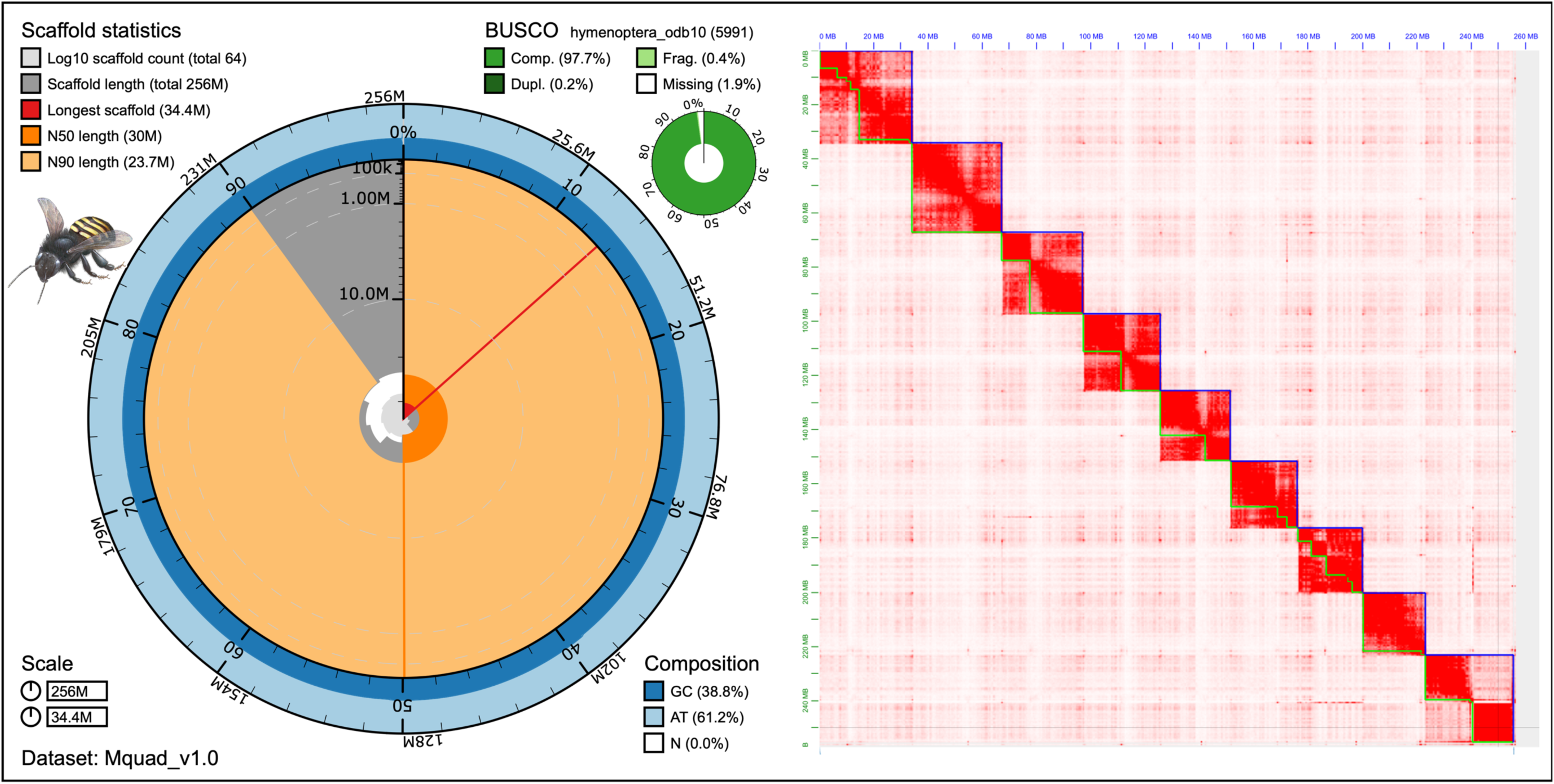
*Melipona quadrifasciata* final genome assembly. A summary representation of the assembly is shown on the left, in which the snailplot is divided into 1,000 size-ordered bins, each representing 0.1% of the total assembly size. Sequence length distribution is shown in dark grey, scaled by the largest scaffold shown in red. Orange and pale orange indicate the N50 and N90, respectively. The pale grey spiral shows the cumulative sequence count on a log scale, with white lines indicating successive orders of magnitude. The blue and pale blue area shows GC and AT distributions. A summary of complete, fragmented, duplicated, and missing BUSCO genes in the hymenoptera_odb10 set is shown in the top right. On the right, the Hi-C contact map of the assembly is shown. Blue lines represent scaffold delimitations, and green lines represent contigs. *M. quadrifasciata* drawing from Birgitte Tümler.

**Figure 2.**
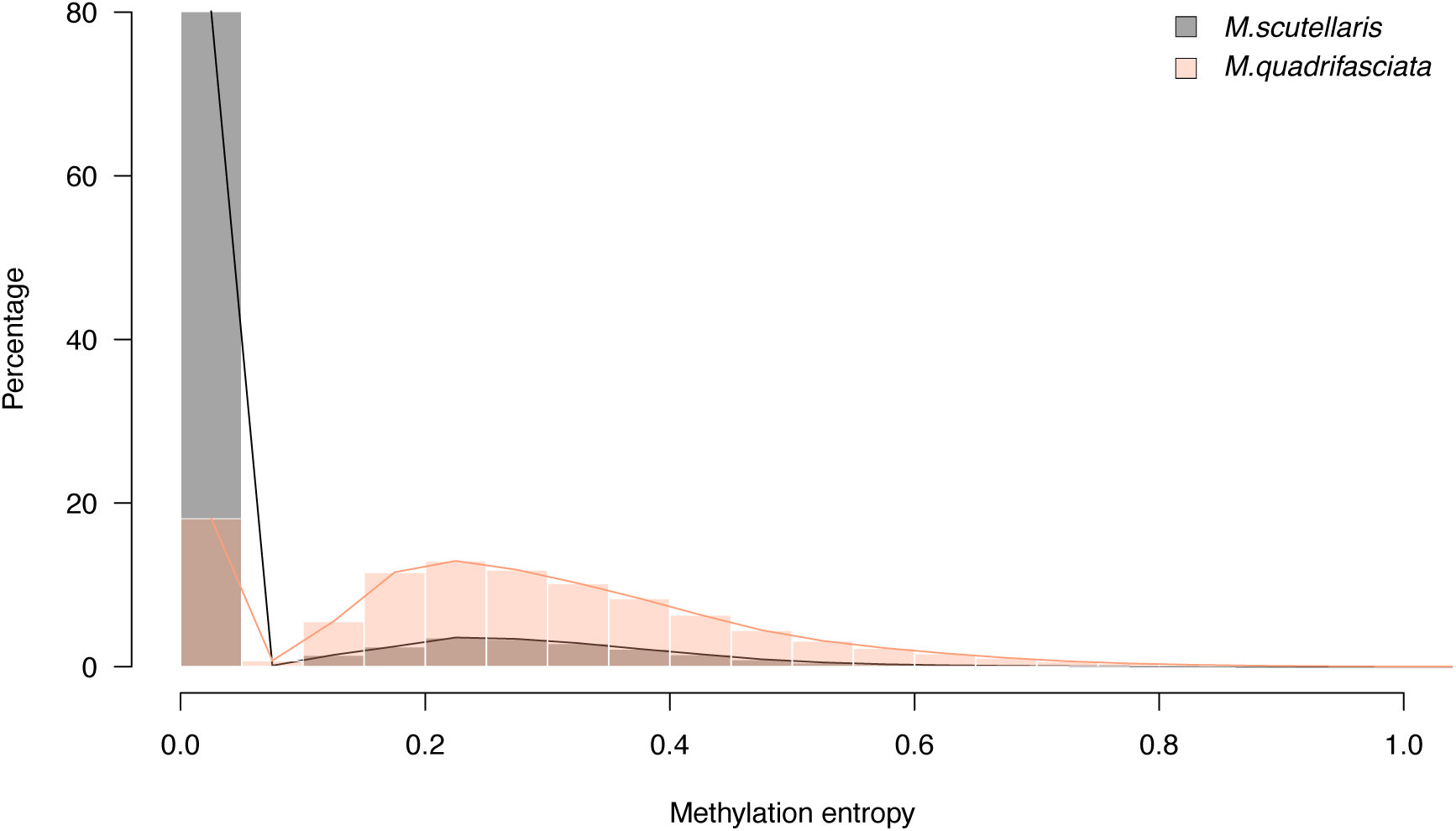
Distribution of methylation entropy at CpG sites across the genomes of *M. quadrifasciata* (salmon bars and line) and *M. scutellaris* (grey bars and black line), estimated on 50bp windows.

**Table I.**
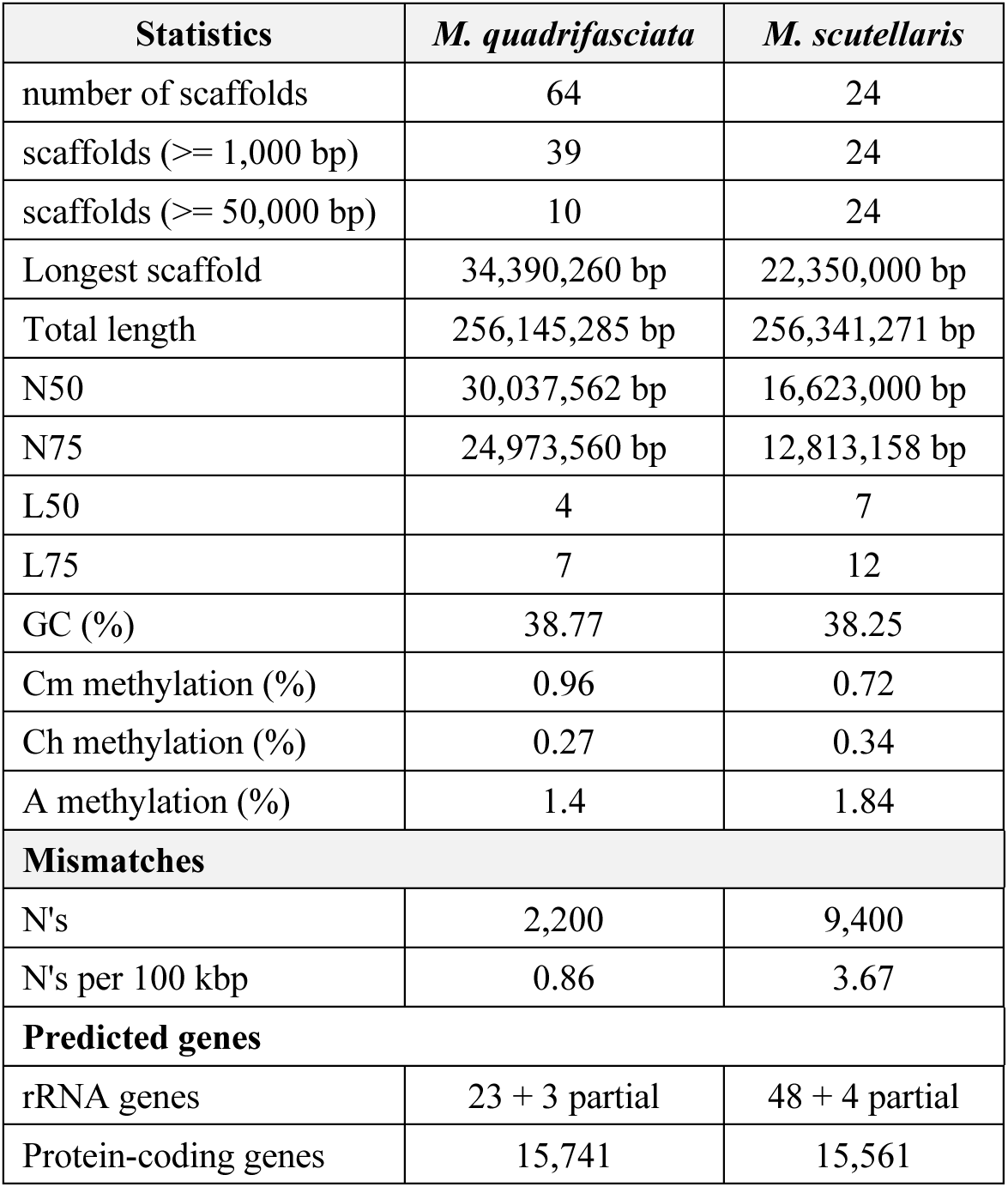
Summary statistics from the final genome assemblies of *M. quadrifasciata* and *M. scutellaris*.

### The genome assembly of *Melipona scutellaris*, over 256 Mb of data in only 24 scaffolds

The *M. scutellaris* genome assembly consisted of 106 contigs totaling about 256.33 Mb before scaffolding, with an N50 of 15 Mb, an L50 of 6, and a maximum contig length of 33,973,957 bp. This dataset returned one hit against the spider *Meta bourneti* (Tetragnathidae) (Figure S1). The contig matching the spider genome was 134,745 bp in length (35.9% mean GC content, 34.59 times of read coverage). Despite matching a non-bee genome, this contig was retained in the assembly because the Tetragnathidae family is of exclusive European distribution (Henriques et al. 2022), and it was unlikely to be a contaminant in our data from which the DNA was sampled and extracted in Brazil. Additionally, the GC content and coverage of the referred contig were consistent with those of the rest of the bee genome. Other contigs matched the Apidae (256,073,566 bp) or the Megachilidae (123,560 bp) families. In total, 131,216,934 paired reads were sequenced from the Hi-C library, and 916,441 possible restriction sites (GATC) were identified in the genome of *M. scutellaris*. Despite similar sequencing depths between the Hi-C data of *M. scutellaris* and *M. quadrifasciata*, the number of high-quality mapping reads retained was smaller for *M. scutellaris* after alignment-based quality trimming, yielding 38,583,392 paired reads (∼21 times the genome coverage). Following Hi-C conformation scaffolding, the assembly initially contained 28 scaffolds, which were refined to 25 after manually correcting one misjoin and three translocations. Long-read realignment across the final genome assembly resulted in ∼105x mean read coverage (SD=1,945) with a mean mapping quality of 58.1.

After removing the mitochondrial genome contig, the final nuclear genome assembly of *M. scutellaris* includes 24 scaffolds spanning 256,341,271 bp (Figure 3). The N50 of this assembly is 16,623,000 bp (about half that of *M. quadrifasciata*), with an L90 on scaffold 15 and a GC content of 38.25% (Table I). Unlike *M. quadrifasciata*, the nine largest scaffolds do not represent complete pseudo-chromosomes in *M. scutellaris* (Figure 3). Instead, the largest 18 scaffolds contain over 99% of the assembly (Figure S2), suggesting that centromeric regions were not fully resolved, potentially due to differing chromosomal domain arrangements between the two *Melipona* species (Pompolo et al. 2002). BUSCO analysis revealed 97.5% completeness of core orthologs in the genome (Figure 3). Possible NUMTs were identified within two scaffolds (Figure S3), with alignments in regions ranging from 1,138 to 11,521 bp, including one large NUMT spanning 65,7% of the mitochondrial genome. Altogether 52 rRNA genes (48 complete and four partial sequences) were identified (Table I).

**Figure 3.**
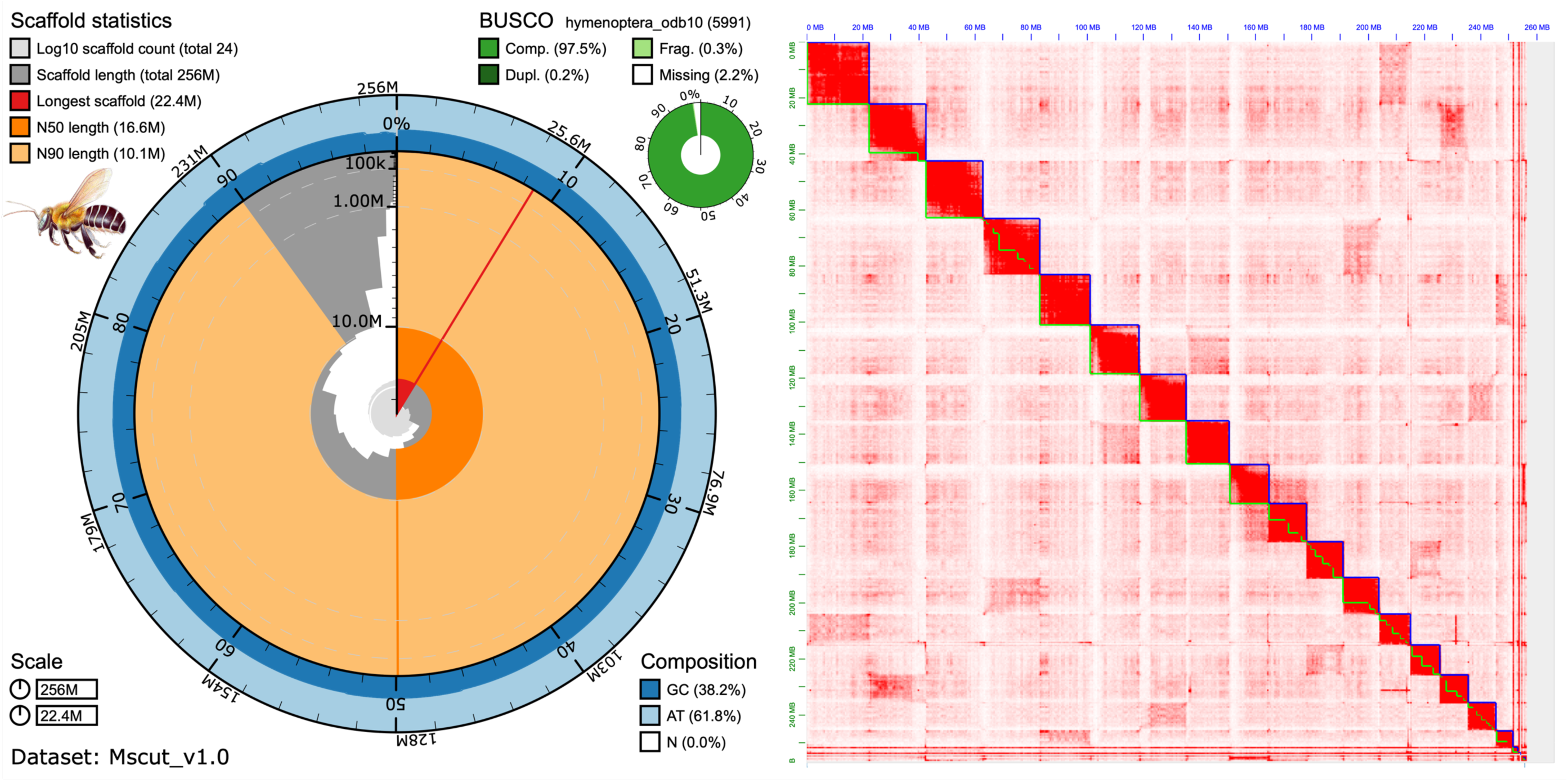
*Melipona scutellaris* final genome assembly. A summary representation of the assembly is shown on the left, in which the snailplot is divided into 1,000 size-ordered bins, each representing 0.1% of the total assembly size. Sequence length distribution is shown in dark grey, scaled by the largest scaffold shown in red. Orange and pale orange indicate the N50 and N90, respectively. The pale grey spiral shows the cumulative sequence count on a log scale, with white lines indicating successive orders of magnitude. The blue and pale blue area shows GC and AT distributions. A summary of complete, fragmented, duplicated, and missing BUSCO genes in the hymenoptera_odb10 set is shown in the top right. On the right, the Hi-C contact map of the assembly is shown. Blue lines represent scaffold delimitations, and green lines represent contigs. *M. scutellaris* drawing from Birgitte Tümler.

We predicted the genome of *M. scutellaris* to comprise 15,561 protein-coding genes, of which 91.6% (14,257) returned significant matches in functional annotation databases, including 12,235 Pfam matches. The predicted proteins reached a BUSCO completeness of 97.6% (C=97.6% [S=77.4%, D=20.2%], F=0.7%, M=1.7%) and comprised 13,835 transcripts with a mono-to-multi-exonic transcript ratio of 0.13 (monoexonic transcripts = 1,569 and multiexonic transcripts = 12,266). Compared to other Aculeata proteomes, the genome annotations of both *M. quadrifasciata* and *M. scutellaris* show higher numbers of protein-coding genes and gene duplications (Figure S4). This is likely justified by the fact that our annotations also have fewer unknown proteins, a notable improvement for stingless bee genomes. The estimated mean methylation for *M. scutellaris* was 0.72% for Cm, 0.34% for Ch (pass threshold ≥ 0.74609375), and 1.84% for A (pass threshold ≥ 0.7714844). DNA methylation entropy at CpG sites in one female worker pupa is shown in Figure 2, and it shows that most sites have zero entropy. This indicates that methylation status is highly conserved across cell tissues in *M. scutellaris*, unlike *M. quadrifasciata*.

### The proportion of repetitive elements is conserved in the genome of the two bees, but the relative abundance of retroelements and DNA transposons differs

Repetitive sequences accounted for a similar proportion of the genomes of *M. quadrifasciata* and *M. scutellaris*, with 18.50% (47,383,871 bp) and 18.75% (48,072,153 bp) of their genomes masked for repetitive elements, respectively (Table II). In both species, most repeats were unclassified, representing 10.58% and 11.72% of the genomes in *M. quadrifasciata* and *M. scutellaris*, respectively. Rolling-circle elements, simple repeats, and low-complexity sequences were also detected at comparable levels in the two species. However, notable differences were observed in retroelements and DNA transposon distributions. In *M. quadrifasciata*, DNA transposons constituted the largest fraction of transposable elements (3.64% DNA transposons vs. 1.18% retroelements), whereas in *M. scutellaris*, retroelements were relatively more abundant (2.22% retroelements vs. 1.78% DNA transposons). All retroelement categories were present at higher frequencies in *M. scutellaris*, with long terminal repeat (LTR) elements showing the most substantial increase (∼1.8-fold more frequent). Specifically, Gypsy/DIRS1 LTR elements were the most abundant retroelements in this category, with a 2.74-fold increase in *M. scutellaris* when compared to *M. quadrifasciata*. Furthermore, two retroviral transposable elements were identified in *M. scutellaris*, while none were detected in *M. quadrifasciata*. Conversely, DNA transposons were approximately twice as abundant in *M. quadrifasciata* than in *M. scutellaris*, and this difference was primarily driven by the greater abundance of hobo-Activator elements (∼2.44-fold higher in *M. quadrifasciata*) and Tc1-IS630-Pogo elements (∼2.54-fold higher in *M. quadrifasciata*).

**Table II.**
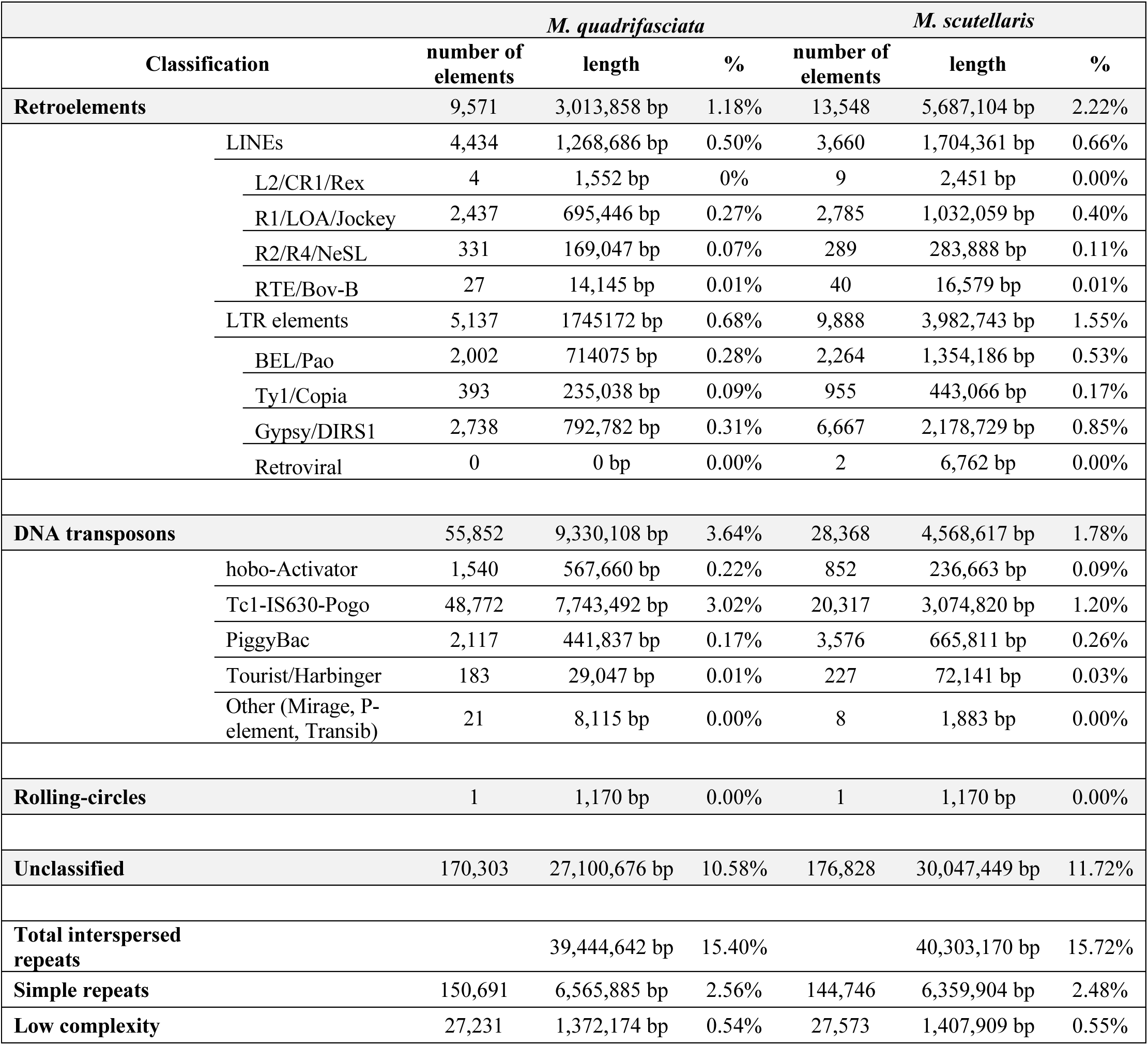
Summary report of all repeats masked in the genomes of *M. quadrifasciata* and *M. scutellaris*.

### Genomic synteny between the two *Melipona* species is mostly conserved

The overall genomic synteny between *M. quadrifasciata* and *M. scutellaris* is largely conserved (Figure 4A). Scaffolds 1, 4, 5, and 6 in *M. quadrifasciata* exhibit no structural differences from their counterparts in *M. scutellaris* (Figures S5, S8, S9, and S10). Shuffling events in scaffolds 2, 3, and 9 of *M. quadrifasciata* are predominantly associated with terminal or centromeric regions. The most notable structural difference between the two genomes occurs in scaffold 7 of *M. quadrifasciata* and the corresponding scaffolds (5 and 17) in *M. scutellaris*. A detailed synteny analysis of these scaffolds (Figure 5) shows that shuffling events in this region were associated with TE accumulation and alterations in GC patterns. Indeed, differences in TE distribution across all scaffolds (Figures S5 to S12) are the most distinct features between the synteny of the two *Melipona* species. *M. scutellaris* shows a higher accumulation of retroelements in several regions of the genome, while the corresponding regions in *M. quadrifasciata* are enriched with DNA transposons. This pattern suggests that the variation in TE ratios between the two species may be driven by an accumulation of retroelements in *M. scutellaris* at sites that harbor DNA transposons in *M. quadrifasciata*.

**Figure 4.**
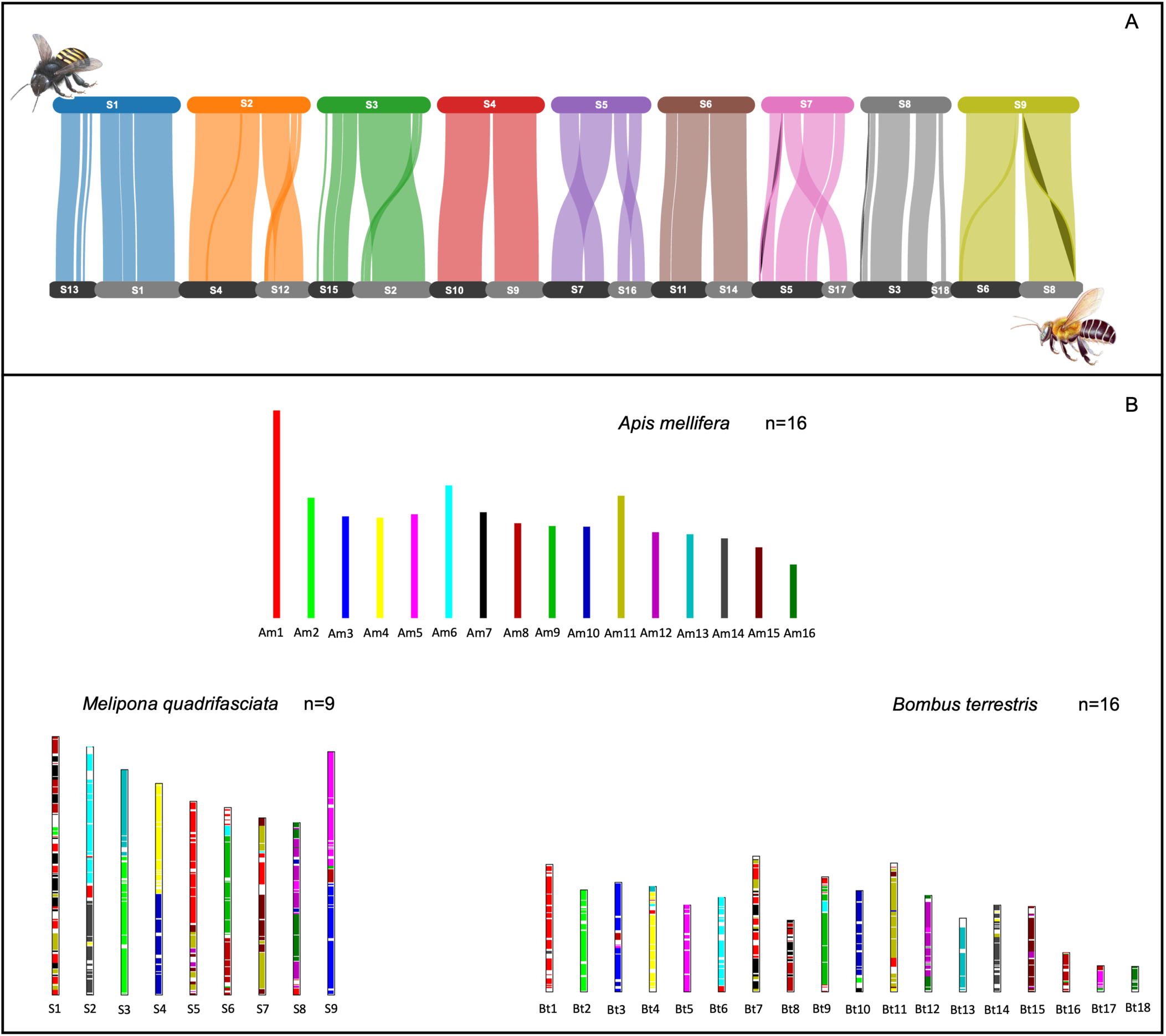
Syntenic regions between bee genomes based on the alignments of the proteomes. **A**-between *M. quadrifasciata* (colored upper bars) and *M. scutellaris* (grey bottom bars) scaffolds. **B**-Between *Apis mellifera* (top) and *Melipona quadrifasciata* (left), and *A. mellifera* and *Bombus terrestris* (right). In B, colors represent regions in *Apis mellifera* chromosomes. Bee drawings from Birgitte Tümler.

**Figure 5.**
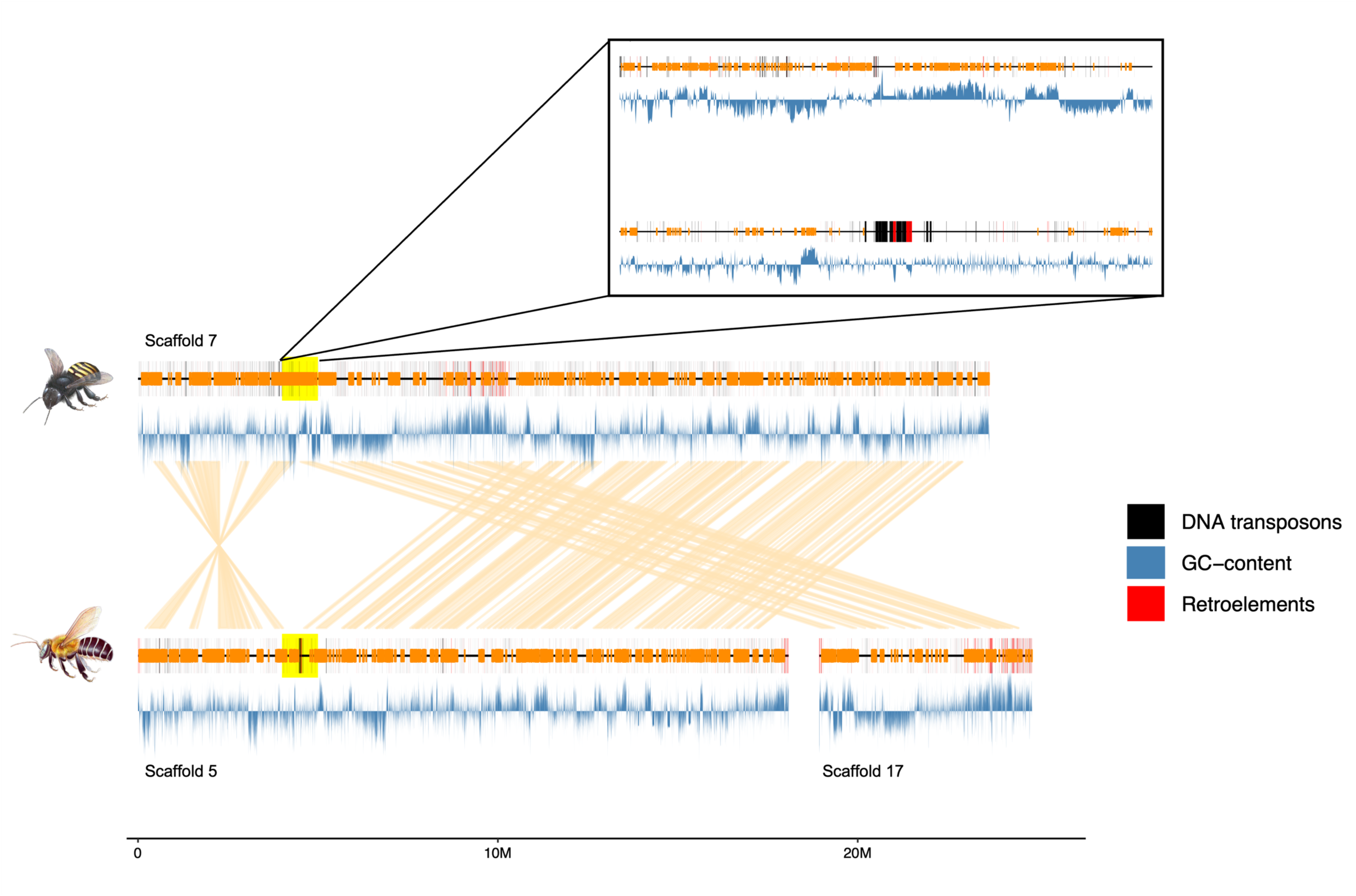
Synteny analysis between *M. quadrifasciata* scaffold 7 and *M. scutellaris* corresponding scaffolds (5 and 17). The region involved in inversion and rearrangement is highlighted in yellow, with a zoomed-in view provided in the inset. Orange blocks represent annotated genes along the scaffolds, while orange lines indicate syntenic connections between genes in the two species. The GC content is depicted as a blue line across the scaffolds, and repetitive elements are marked: retroelements in red and DNA transposons in black.

When comparing synteny across all eusocial corbiculate bees (Figure 4B), the analyses confirm that the reduction in chromosome number observed in the genus *Melipona* is primarily a result of chromosomal rearrangements and fusions, as previously proposed (Araujo et al. 2024). For example, scaffold 1 in *M. quadrifasciata* is syntenic to multiple chromosomes in *A. mellifera* (Am1, Am2, Am7, Am8, and Am11), reflecting extensive shuffling events that occurred during the divergence of these lineages. However, many of these rearrangements are shared between the *Melipona* and two chromosomes in the bumble bee *Bombus terrestris* (Bt7 and Bt8), supporting the conclusion that this structure is a synapomorphic feature of their common ancestor. Given that n=18 is the most likely haploid chromosome number of this corbiculate ancestor (Travenzoli et al. 2019), it is plausible that a further fusion event in *Melipona* resulted in the large scaffold 1. Scaffolds 3 and 4 in *M. quadrifasciata* exhibit more conserved synteny across all bee lineages, with only chromosomal fusions observed in *Melipona*. For example, scaffold 3 is a fusion from Am2 and Am13 of *A. mellifera* (or Bt2 and Bt13 of *B. terrestris*).

### Structural variants between the two *Melipona* species coincide with TE accumulation and scaffold edges

The graph-based chromosome alignment using Minigraph-Cactus revealed extensive structural variation between the genomes of *M. quadrifasciata* and *M. scutellaris*. Across all chromosomes, we identified a total of 59,252 structural variants (≥50 bp), comprising 24,666 insertions (41.6%) and 34,586 deletions (58.4%). The sizes of SVs ranged from 50 to 137,004 bp, with a median size of 87 bp (Table SI).

Quantification of SVs within 100-kb windows revealed a non-uniform distribution of SVs across the genome, with a pronounced accumulation of insertions near the edges of each *M. scutellaris* scaffold (Figure 6). We also observed a negative correlation between the number of deletions and the number of genes across many regions in the genome, including scaffold 6, scaffold 7, and scaffold 8. Clusters of SVs also coincided with areas of high TE density. For instance, a cluster of insertions on scaffold 8 overlapped with a region of low gene frequency but high TE accumulation (Figure 6).

**Figure 6.**
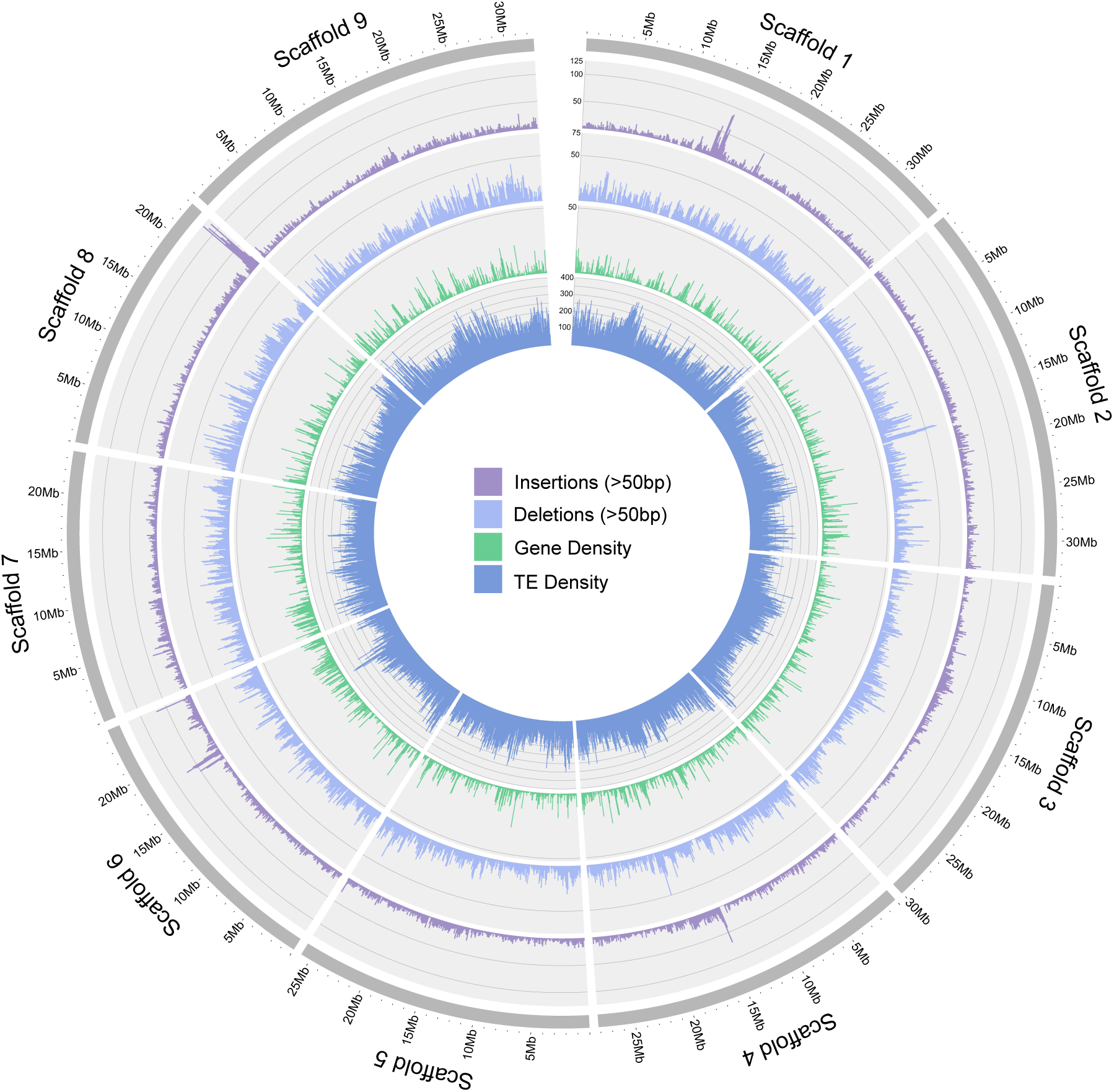
Circos plot showing the distribution of SVs on nine pseudo-chromosomes utilizing *M. quadrifasciata* as the reference in each of the following SV categories: insertions (>50bp), deletions (>50bp), gene density, and TE density. Insertions are segments exclusive to *M. scutellaris*, while deletions are missing sequences in this species.

### Gene content evolution shows that *Melipona* Group II species have more histone deacetylase orthologs than Group I

We constructed a genomic dataset matrix comprising 284 single-copy orthologs. During tree estimation, these were divided into 34 partitions, with an average of 8.3 orthologs per partition, spanning 101,094 sites. Among these, 30,081 (29.7%) were parsimony-informative, while the remaining were constant (n = 49,708, 48,2%) or singletons (n = 21,305, 21,1%). The resulting phylogenomic tree (Figure S13) displayed robust support across all clades (UFBoot >= 98). As anticipated, stingless bees and bumble bees were resolved as sister groups, diverging in our phylogeny, approximately 53.3 Mya during the early Miocene. Within stingless bees, *Melipona* diversification occurred around 8.2 Mya in the late Miocene. In our analyses, *M. quadrifasciata* was the basal taxon of the *Melipona* clade.

We identified 399,846 orthologs, 378,945 of which clustered into 15,158 orthogroups, achieving an assignment rate of 94.8%. Nearly 40% of the orthogroups (n=6,065) were shared across all species, while roughly 15% (n=2,283) were species-specific. *Melipona quadrifasciata* and *M. scutellaris* showed comparable numbers of orthologs but differed in species-specific orthogroups: 11 species-specific orthogroups were reported in the former (encompassing 26 genes) and 18 in the latter (encompassing 46 genes) (Table SII). Gene family content evolution analysis identified 143,309 size change events (λ = 0.004166), with 1,346 orthogroups (8.8%) showing significant changes (p < 0.05). Contractions (n = 94,358, 65.8%) occurred nearly twice as often as expansions (n = 48,951, 34.2%). In *M. quadrifasciata*, we identified 1,939 contractions and 1,091 expansions, while *M. scutellaris* showed 1,814 contractions and 1,135 expansions (Table SIII). Additionally, 2,465 rapid evolution events across 654 orthogroups were identified, including 1,511 expansions (61.3%) and 954 reductions (38.7%).

In total, 193 rapid evolution events were species-specific either to *M. quadrifasciata* (n=92) or *M. scutellaris* (n=91). Among these, the most frequent genes were zinc finger proteins (6 genes; 2 in *M. quadrifasciata* and 4 in *M. scutellaris*), odorant receptors (5 genes containing a 7tm_6 Pfam motif; 3 in *M. quadrifasciata* and 2 in *M. scutellaris*), cytochrome P450 (4 genes; two in each species), and guanylate kinase homologous (3 genes; 1 in *M. quadrifasciata* and 2 in *M. scutellaris*) (Supplementary file 2). In both species, the species-specific genes under rapid evolution were mostly associated with terms involved in development (e.g., GO0001748 insect visual primordium development, GO:0002164 larval development, GO:0009888 tissue development, and GO:0021700 developmental maturation), metabolism (e.g., GO:0006518 peptide metabolic process, GO:0043170 macromolecule metabolic process, GO:0006259 DNA metabolic process), environmental response (e.g., GO:0009629 response to gravity, GO:0006955 immune response, GO:0009605 response to external stimulus), regulation of cellular processes (e.g., GO:0046782 regulation of viral transcription, GO:0010646 regulation of cell communication, GO:0019222 regulation of metabolic process), and cellular organization (e.g., GO:0043954 cellular component maintenance, GO:0006325 chromatin organization, GO:0043933 protein-containing complex organization) (S2). These findings underscore the role of these biological functions in lineage-specific adaptations of stingless bees.

A notable finding was the considerable variation in histone deacetylase-related orthologs among the stingless bee genomes, ranging from 11 genes in *M. quadrifasciata* to 37 in *Frieseomelitta varia* (Table SIV). These enzymes catalyze acetyl group removal, enhancing DNA-histone binding and promoting heterochromatin formation (Park and Kim 2020). *Melipona scutellaris* (Group II) exhibited 23 histone deacetylase orthologs, significantly more than both species of Group I, *M. quadrifasciata* (n = 11) and *M. bicolor* (n = 12) (F = 3.5, p = 0.03999). The counts of other chromatin-modifying orthologs, including acetyltransferases, demethylases, and methyltransferases, showed no significant variation across stingless bees.

## Discussion

This study provides high-quality, pseudo-chromosome-level genome assemblies for two *Melipona* species with diverging heterochromatin organization: *M. quadrifasciata* and *M. scutellaris*. Despite their conserved genome sizes and genomic synteny, these species exhibit notable differences in transposable element distribution, structural variants, and epigenetic landscapes. *M. scutellaris* shows a pronounced accumulation of retroelements (particularly Gypsy/DIRS1) linked to structural variants, histone deacetylase-containing orthogroups expansion, and alterations in genome methylation entropy patterns compared to *M. quadrifasciata*. These genomic and epigenetic differences likely contribute to the diverging heterochromatin organization observed in these species, offering insights into the evolutionary forces shaping heterochromatin and chromosomal evolution in eusocial insects.

The genome of *M. quadrifasciata* was the first to be published for a stingless bee (NCBI ASM127656v1) (Kapheim et al. 2015). While groundbreaking and adequate in size (∼256.3 Mb), this assembly was highly fragmented (2,866 scaffolds; scaffold N50=1.9 Mb) due to its reliance on short-read sequencing data only. Consequently, it contained more unknown and missing orthologs (Figure S4). In contrast, our newly sequenced and assembled genome of *M. quadrifasciata* spans a similar length (∼256.14 Mb) but with over 1,000 additional predicted genes (15,741 vs. 14,711), all distributed across only 64 scaffolds, with the largest nine representing pseudo-chromosomes. Similarly, our unprecedented *M. scutellaris* genome assembly is an important resource for both the scientific and stingless bee-keeping communities, as this species is amongst the most broadly studied and reared. Indeed, the two genomes produced are the most contiguous for stingless bees to date, and the sole ones supported by Hi-C data for reaching chromosome-level scaffolds.

These genomes offer a unique opportunity to explore heterochromatin dynamics in genomic evolution. In *Melipona*, karyotypic analyses of several lineages have consistently shown a bimodal heterochromatin distribution across species (Pompolo et al. 2002; da Cunha et al. 2018). Group I species, like *M. quadrifasciata*, typically exhibit less than half of their karyotypes marked by C-banding, predominantly in pericentromeric regions, indicating lower levels of heterochromatin. Conversely, Group II species, such as *M. scutellaris*, display over 50% of their karyotypes marked by C-banding, reflecting high heterochromatin levels (Pompolo et al. 2002). Species from Group I are from the subgenera *Eomelipona*, *Melikerria*, and *Melipona sensu stricto*, while Group II species belong to the *Melikerria* and *Michmelia* subgenera (Pereira et al. 2021). The subgenus *Melipona sensu stricto*, which includes *M. quadrifasciata*, shows the earliest divergence among these four subgenera (Lepeco et al. 2024), suggesting that reduced heterochromatin represents the ancestral state of the group. Further cytogenetic evidence supports at least two independent origins of high heterochromatin levels within the clade, one in *Melikerria* and one in *Michmelia* (Cunha et al. 2020). These events were hypothesized to be driven by repetitive element expansion (Cunha et al. 2020), which should, in turn, lead to an augmentation of the genome size in species of Group II (Pereira et al. 2021; Tavares et al. 2010).

Our findings corroborate the divergence in chromosomal organization. Experimental challenges during DNA sequencing and Hi-C library preparation revealed intrinsic genomic differences between the two *Melipona* species. For instance, the sequencing of *M. quadrifasciata* was more efficient, requiring data from a single flow cell, whereas *M. scutellaris* required multiple flow cells and flow cell washes due to a rapid decline in pore availability. The Hi-C data for *M. scutellaris* also yielded about half the output coverage compared to *M. quadrifasciata*. These challenges are unlikely to result from experimental differences as, in both cases, sampling of the bees, DNA extraction, and library preparation were performed by the same researcher and protocols. However, we did not find evidence to support the hypothesis that these differences result from an augmentation of repetitive elements and genome sizes. Both species exhibit highly conserved genome sizes (256.14 Mb for *M. quadrifasciata* and 256.34 Mb for *M. scutellaris*), with comparable repetitive element ratios (∼18.5%). Instead, the key differences lay in their TE composition.

In *M. scutellaris*, a marked expansion of Gypsy/DIRS1 retroelements, coupled with the presence of retroviral sequences, has led to a greater retroelement complement compared to *M. quadrifasciata*, which instead harbors a higher proportion of DNA transposons. This dynamic turnover has likely driven structural rearrangements that may have contributed to the chromosomal evolution of these species. This assumption is supported by the association of retroelements with chromosomal alterations in the synteny analyses and by the higher frequencies of SVs in regions of elevated TE density found in *M. scutellaris*. These findings are consistent with previous studies on insects, suggesting that TEs might act as major drivers of genome reorganization. For instance, in *Drosophila melanogaster*, spontaneous TE insertions have caused over half of all the known phenotypic mutations (González et al. 2008). Furthermore, retrotransposon expansions by a cut-and-paste mechanism are known to induce SVs, alter gene expression, and rearrange chromosomes (Kabi and Filion 2021; Liao et al. 2023). Our structural variation analysis revealed a negative correlation between SV deletions and gene number reduction, underscoring the selective pressures associated with TE activity.

Heterochromatin and DNA methylation are key epigenetic mechanisms that suppress TE and endogenous retrovirus activity (Kabi and Filion 2021). Our data show that in *M. scutellaris*, TE dynamics are accompanied by distinct epigenetic patterns, including reduced methylation entropy, compared to *M. quadrifasciata*. Methylation entropy measures the randomness of DNA methylation within a sample (Xie et al. 2011). High entropy indicates that sequenced reads exhibit methylated cytosines at random positions, while zero entropy signifies a conserved methylation signal, in which all cytosines at the same position display consistent methylation status across sequenced cells (Xie et al. 2011). We used whole bodies for methylation data of both species, still, most methylated sites in *M. scutellaris* exhibited zero entropy, indicating highly conserved methylation patterns across cell types, contrasting with the more variable methylation entropy observed in *M. quadrifasciata* (Figure 2). This epigenetic signature may reflect an active regulation of retroelements conserved across different tissues, potentially mitigating their mutagenic effects while maintaining genome stability. The expansion of histone deacetylase-associated orthologous in *M. scutellaris* further suggests a role for histone modification in heterochromatin regulation, providing a mechanistic explanation underlying the high heterochromatin levels observed. These findings indicate that epigenetic mechanisms in *M. scutellaris* may have co-evolved with retrotransposon activity to preserve chromosome integrity, ultimately contributing to the distinct heterochromatic pattern characteristic of Group II species.

While this study advances our understanding of genome evolution in stingless bees, especially in the species-rich genus *Melipona*, further exploration is needed. Expanding genomic analyses to additional *Melipona* species would clarify the evolutionary trajectories of chromosomal architecture and heterochromatin organization in all *Melipona* subgenera. Furthermore, detailed transcriptomics and epigenomics studies across developmental stages and castes could further elucidate functional implications of the TE dynamics and heterochromatin variation exemplified here to the biology and ecology of the highly diverse but underexplored stingless bees.

## Conclusions

This study delivers the first high-quality, pseudo-chromosome-level genome assemblies for two *Melipona* species, providing valuable resources for understanding the genomic and epigenetic dynamics in the evolution of the highly eusocial stingless bees (Meliponini). By comparing the genomes of *M. quadrifasciata* and *M. scutellaris*, we uncovered significant differences in transposable element composition, structural variation, and epigenetic factors, which likely contributed to their distinct heterochromatin organizations. Our findings indicate that changes in TE dynamics may affect the genomic organization without leading to genome size expansion. Beyond offering insights into genome evolution in the genus *Melipona*, these data pave the way for future research, fostering a deeper understanding of genome architecture and its ecological and evolutionary consequences in these key pollinators.

## Methods

### Data generation

We sampled female pupae from colonies of *M. quadrifasciata* and *M. scutellaris* by opening the brood and checking the appropriate developmental stage. Upon collection, all bees were immediately frozen in liquid nitrogen and stored at -80 °C until processing. The colonies used in this study were maintained at the University of São Paulo, Ribeirão Preto Campus (Brazil). Total DNA and RNA for long-read and RNA-Seq sequencings were extracted *in situ* from the whole body of pupae following the Qiagen AllPrep DNA/RNA/miRNA Universal Kit protocol. DNA conformation data was extracted and processed from a single pupa per species with the Arima High Coverage HiC Kit (November 2021 version) and the Library Preparation for Arima High Coverage HiC Kit (May 2023 version) protocols. Genome sequencing was conducted using Oxford Nanopore technology (Mk1C device) with flow cells version 10.4.1 and the LSK114 library preparation kit, following the manufacturer’s instructions. Long-read basecalling was performed with Dorado v0.3.2 ([CSL STYLE ERROR: reference with no printed form.]) using the duplex superior basecalling mode (dna_r10.4.1_e8.2_400bps_sup@v4.2.0). RNASeq library preparation and sequencing of 100 bp paired reads were performed by Macrogen (South Korea facility) using the TruSeq Stranded mRNA Library Prep Kit and the Illumina platform. The Arima libraries were sequenced on the Illumina NovaSeq at the GIGA Institute, University of Liège (Belgium) to generate 60 million 150 bp paired-end reads per library.

Long reads were also used for DNA methylation estimations. Methylation calls were carried out with Dorado in simplex mode to detect 5mC and 5hmC methylation (using the sup,5mC_5hmC options). Summary mean methylation (based on 10% of the total data) and methylation entropy of the reads were assessed with modkit v0.3.1 ([CSL STYLE ERROR: reference with no printed form.]). Methylation analyses were based on the sequencing data generated from the whole body of a single female worker pupa from each species.

### Genome assembly

Quality assessments of the reads, both before and after trimming, were performed using FASTQC v0.12.1 (Andrews 2010) and QUAST v5.0.2 (Gurevich et al. 2013). We filtered reads with low-quality scores (phred score < 10) and removed the first 20 nucleotides (due to bias and reduced quality) using Nanofilt v2.8.0 (De Coster et al. 2018) (parameters: -q 10, -- headcrop 20). For *M. quadrifasciata,* we generated sequencing data from a single worker bee on one flow cell, resulting in 5,990,855 trimmed long reads (43 Gb), with an N50 of 7,947 bp, a maximum read length of 150,719 bp, and estimated coverage of 90x. For *M. scutellaris,* sequencing data from four workers of the same colony (each sequenced on a separate flow cell) were combined to yield 4,164,074 reads (49 Gb) with an N50 of 12,987 bp and a maximum read length of 298,388 bp. Based on preliminary assembly tests, we removed mitochondrial reads from these datasets to avoid possible NUMTs interference. For this, before the nuclear genome assembly, the mitochondrial genomes of both species were assembled with MitoHifi v2 (Uliano-Silva et al. 2023) using the *M. bicolor* mitochondrial genome (AF466146 (Araujo and Arias 2019)) as the reference for *M. quadrifasciata* and the mitochondrial genome of *M. scutellaris* (accession NC_026198 (Pereira et al. 2016)) as reference for the same species. The assembled mitochondrial contigs were then used as bait to remove the mitochondrial reads. We aligned all trimmed reads to the species’ respective mitochondrial contig using minimap2 v2.22 (Li 2018) (alignment options: --secondary=no -ax map-ont) and removed the aligned reads from the datasets with a custom script [https://github.com/nat2bee/fasta-manipulation/blob/master/FastaChomper.py].

The two *Melipona* species required distinct genome assembly strategies. For *M. quadrifasciata*, we used Flye v2.9.1 (Kolmogorov et al. 2019) (parameters: -asm-coverage 90 -g 250m -no-alt-contigs -t 30 -scaffold) followed by polishing with Medaka v1.7.2 ([CSL STYLE ERROR: reference with no printed form.]) (parameter: -m r1041_e82_400bps_sup_g615). The genome of *M. scutellaris* was assembled by merging two initial assemblies: one generated with Flye (as described for *M. quadrifasciata*) and another by NextDenovo v2.5.2 (Hu et al. 2024) (parameters: read_cutoff = 1k, genome_size = 250m, sort_options = -m 40g -t 15). Flye produced an assembly with a high BUSCO completeness score but highly fragmented, while NextDenovo produced a more contiguous assembly with lower BUSCO completeness. Both assemblies were polished with Medaka and then merged with quickmerge (Solares et al. 2018) (parameter: -l 2150149, using the Flye assembly as the hybrid assembly and the NextDenovo results as the self-assembly). The resulting merged assembly was subsequently polished once again with Medaka. As the assemblies of both species were generated from diploid females, we used purge_dups v1.2.5 ([CSL STYLE ERROR: reference with no printed form.]) with the recommended cutoffs to remove potential haplotigs. To screen for possible contaminants, we used the BlobToolKit v4.1.5 (Challis et al. 2020) pipeline, comparing our assemblies to the NCBI (using blastn, blast+ v2.12.0 (Camacho et al. 2009)) and the Uniprot (with diamond blastx, v2.1.8 (Buchfink et al. 2021)) databases. Search parameters for blastn were -db nt -outfmt 6 qseqid staxids bitscore std -max_target_seqs 10 -max_hsps 1 -evalue 1e-25, and for diamond, -outfmt 6 qseqid staxids bitscore qseqid sseqid pident length mismatch gapopen qstart qend sstart send evalue bitscore -sensitive -max-target-seqs 1 -evalue 1e-25. Coverage of the contigs was evaluated in BlobToolKit by realigning the reads to the genomes using minimap2 (parameter: *-ax map-ont*).

For further scaffolding, we generate over 60 million 150 bp paired-end reads per species from each 3D conformation Hi-C library. The Hi-C sequencing data were aligned to the assemblies using BWA v0.7.17 (Li 2013) (alignment options: mem -A1 -B4 -E50 -L0). Then, with HiCExplorer tools v3.7.2 (Wolff et al. 2020), we identified restriction sites and filtered valid paired reads for matrix building (parameters: binSize 10000 --restrictionSequence GATC -danglingSequence GATC -inputBufferSize 400000). Scaffolding was performed with YaHS v1.2 (Zhou et al. 2023) based on the filtered alignments, using the Juicer v1.6 (Durand et al. 2016b) and Juicer Tools v2.20 (Durand et al. 2016a) implementations. Manual curation of scaffolded assemblies was conducted in Juicebox v2.15 (Durand et al. 2016a).

The quality and statistics of the final assemblies were assessed using QUALIMAP v2.3 (García-Alcalde et al. 2012), BUSCO v5.1.2 (Simão et al. 2015) (hymenoptera_odb10, 2024-01-08 database), OMArk web browser v0.3.0 (OMAmer database: OMA from November 2022, and LUCA, OMAmer version: 2.0.0) (Nevers et al. 2022), and QUAST v5.0.2 (Gurevich et al. 2013). Potential mitochondrial NUMTs in the nuclear genome were identified by aligning the mitochondrial genome to the final nuclear assembly using LAST v1522 (Kiełbasa et al. 2011). We used the default scoring and mismatch probability threshold >1e-2, applying two-way alignments (i.e., aligned the mitochondrial genome to the nuclear genome and vice versa) to confirm NUMTs.

### Genome annotation

Repetitive regions in the genomes were identified following the protocol outlined at https://github.com/nat2bee/repetitive_elements_pipeline. Briefly, we built custom repeat libraries using RepeatModeler v2.0.5 (Flynn et al. 2020), TransposonPSI (Haas 2010), and LTRharvest from GenomeTools v1.6.5 (Ellinghaus et al. 2008). These libraries were merged into a single non-redundant library using USEARCH v11.0.667 (Edgar 2010) (<80% identity). Repetitive elements in our custom libraries were then annotated with RepeatClassifier v2.05 and combined with the Dfam v3.8 Hymenoptera library included in RepeatMasker v4.1.6 (Smit et al. 2013). The final repetitive element libraries are at https://github.com/nat2bee/repetitive_elements_pipeline/tree/master/database. We used these libraries to mask repetitive regions in the genomes with RepeatMasker.

Protein-coding genes were predicted with the BREAKER3 (Gabriel et al. 2024) pipeline, which integrates species-specific transcript and protein evidence. Protein evidence was sourced from the OrthoDB 11 Arthropoda database. For transcriptomic evidence, we sequenced total RNA from three workers and three queen pupae per species (one individual per colony, totaling three colonies). RNA-Seq reads were aligned to the reference genomes using hisat2 v2.2.1 (parameter: –data) (Kim et al. 2015), and alignments were converted and sorted to bam files using SAMtools v1.16.1 (Li et al. 2009). Functional annotation of the predicted proteins was conducted with eggnog-mapper v2.1.12 (Rodríguez del Río et al. 2023), using the Insecta database (parameters: -m diamond -dbmem -tax_scope Insecta -m hmmer -d Insecta -pfam_realign denovo). The quality and completeness of the gene prediction were assessed using BUSCO in the protein mode and the analyze_exons.py script from the GALBA repository (Gaius-Augustus/GALBA/master/scripts/analyze_exons.py). Ribosomal RNA sequences were predicted using Barrnap v0.8 implemented within QUAST v5.0.2 with the option -k euk.

### Genomic synteny

We assessed synteny between the two *Melipona* genomes and compared the pseudo-chromosomes of *M. quadrifasciata* with those of the two other eusocial corbiculate bee lineages: *Apis mellifera* (Apini) and *Bombus terrestris* (Bombini). To identify the syntenic regions between protein sets, we used blastp from NCBI-blast + v2.12.0 (Camacho et al. 2009) with parameters set to -e 1e-10 -b 5 -v 5, and -m 8. Collinear conserved blocks were then estimated and visualized using MCScanX (Wang et al. 2012). Additional comparative synteny plots were generated with SynVisio (Bandi and Gutwin) and R v4.3.3 (R Core Team 2017), employing the gggenomes and GenomicRanges packages.

### Structural variants

To identify structural variants in *M. quadrifasciata* and *M. scutellaris*, we conducted a graph-based chromosome alignment with Minigraph-Cactus (Hickey et al. 2023). For each chromosome, a separate graph was constructed, utilizing *M. quadrifasciata* (the most contiguous assembly) as the reference and a pair of syntenic *M. scutellaris* scaffolds (Table SV and Figure 4). Large SVs were defined as SVs > 50 bp, where insertions consist of segments present exclusively in *M. scutellaris*, and deletions are segments absent in *M. scutellaris*. SVs were quantified within 100-kb windows in the reference genome and visualized using Circos (Krzywinski et al. 2009).

### Gene content evolution

We identified and clustered orthologous genes into orthogroups using OrthoFinder v.2.5.5 (Emms and Kelly 2019) based on protein data from 19 hymenopterans (Table SVI), including 14 bee species representing the five major families (Andrenidae, Apidae, Colletidae, Halictidae, and Megachilidae) and five non-bee hymenopteran species as outgroups. To assess changes in gene composition within orthogroups, we constructed a phylogenomic tree from the single-copy orthologs identified with BUSCO v.5.4.6 (Simão et al. 2015) and aligned them using MUSCLE v.3.8 (Edgar 2004). Alignments were quality-trimmed with trimAL v.1.4.1 (Capella-Gutiérrez et al. 2009) and then concatenated into a matrix of 90% occupancy using the BUSCO Phylogenomics pipeline (https://github.com/jamiemcg/BUSCO_phylogenomics). This cutoff balanced missing data against the exclusion of phylogenetically informative loci. We inferred a maximum-likelihood phylogenomic tree using IQ-TREE v.2.1.3 (Minh et al. 2020) with the following parameters: -B 1000 (1,000 ultrafast bootstrap replicates), -m MFP- MERGE (model selection with ModelFinder (Kalyaanamoorthy et al. 2017), merging loci until the model fit stops increasing), and --symtest-remove-bad (to filter out loci violating assumptions of stationarity, reversibility, and homogeneity). Before time-calibrating the inferred tree, we generated a control file using r8s v.1.81 (Sanderson 2003) with the following parameters: -s 101094 (total site number in the final multi-loci alignment), -c 240 (calibration age, in million years, for the node (Kumar et al. 2022)). We then manually added two additional calibration points (Aculeata, 118, and Meliponini, 37 (Kumar et al. 2022)) to the control file. Based on the phylogenomic tree and orthogroups identified, we used CAFE v.4.2.1 (De Bie et al. 2006) to detect changes in gene family content across the species with the following parameters: -t 8 -s -t 1 (uniform lambda). The function of fast-evolving orthogroups in the two *Melipona* was based on the annotation of representative genes in their genomes and REVIGO (Supek et al. 2011) was used to visualize the gene ontology biological processes with tiny reduction and the *Drosophila melanogaster* database.

Since we were interested in heterochromatin differences between the two *Melipona* species, we additionally analyzed changes in orthogroups containing histone-interacting enzymes (i.e., acetyltransferases, deacetylases, demethylases, and methyltransferases). For this, orthogroups were functionally annotated based on sequence homology using Diamond v.2.0.14 (Buchfink et al. 2014). First, one representative sequence of each orthogroup (the longest) was aligned against the reviewed Swiss-Prot database, and those without matches were subsequently aligned against the unreviewed TrEMBL database (Bateman et al. 2023). For all eusocial corbiculates, we counted the number of orthologs in orthogroups containing acetyltransferases, deacetylases, demethylases, and methyltransferases. Then, we performed one-way ANOVA statistical tests using the R (R Core Team 2017) package ’stats’ to determine whether the number of orthologs for each specific enzyme differed significantly in the two *Melipona*.

## DATA ACCESS

All data generated for this study are deposited at NCBI under the BioProject PRJNA1206466. SRA accession for long read data from *M. quadrifasciata* is SRP555293, and for *M. scutellaris* are SRR31893452, SRR31893451, SRR31893450, and SRR31893449. RNASeq data accessions are: SRR31893476, SRR31893465, SRR31893458, SRR31893474, SRR31893469, and SRR31893461 for *M. quadrifasciata*; SRR31893467, SRR31893488, SRR31893481, SRR31893490, SRR31893446, and SRR31893484 for *M. scutellaris*. The accession from Hi-C data of M. quadrifasciata is SRR32013089 and for *M. scutellaris* is SRR32013090. Samples used in this study are authorized under the SISBIO project N°81843-1, in collaboration with the Universidade de São Paulo – Brazil, and were exported under the IBAMA permit 23BR045417/DF, respecting the Nagoya agreement.

## ACKNOWLEDGMENTS

We thank Kimberly Walden (*in memorian*) for her valuable insights into SV analysis.

## DECLARATIONS

### Ethics approval and consent to participate

Not applicable.

### Competing interest statement

The authors declare they have no competing interests.

### Funding

This study received funding from the Belgium National Fund for Scientific Research (FRS-FNRS) for NSA (Chargée de Recherches, FC34383) and SA (grant GETME J.0008.23); from the São Paulo Research Foundation (FAPESP) for NSA (visiting research grant 2022/14675-9). RRF thanks Veracel Celulose for the postdoc fellowship (Proc. 23746.001092/2022-30); TMB thanks Suzano Papel e Celulose S/A (grant, Proc. 23746.009802/2021-88).

### Authors’ contributions

All authors contributed to the manuscript preparation and revision, below are listed other specific contributions. NSA: study design, funding, data generation, data analysis, manuscript preparation, and study coordination. PA: study design, data analysis, manuscript preparation. RRF: study design, data analysis, manuscript preparation. LBS: data analysis, manuscript preparation. FR: data generation, manuscript preparation. RCFN: manuscript preparation, and study coordination. MH: data analysis, manuscript preparation, and study coordination. KH: funding, data generation, manuscript preparation, and study coordination. TMB: data analysis, manuscript preparation, and study coordination. SA: study design, funding, manuscript preparation, and study coordination.

## Notes

### Competing Interest Statement

The authors have declared no competing interest.

